# Multi-task fMRI outperforms resting-state fMRI for revealing task-invariant organization of the human brain

**DOI:** 10.64898/2026.03.09.710558

**Authors:** Caroline Nettekoven, Ali Shahbazi, Bassel Arafat, Matea Skenderija, Jinkang Derrick Xiang, Ana Luísa Pinho, Jörn Diedrichsen

## Abstract

Resting-state *functional Magnetic Resonance Imaging* (fMRI) is widely used to infer the intrinsic functional organization of the brain, yet it remains unclear how well this approach can predict the structure of brain activity observed across a diverse set of mental states. Here we compare resting-state to task-based fMRI using diverse task batteries within the same individuals. We find that multi-task fMRI data consistently outperform resting-state estimates in predicting functional organization during novel tasks. This advantage persists across preprocessing strategies, brain regions, and independent datasets. While task activation estimates do show task-dependency when using only few tasks, increasing task diversity reduced task-specific bias, with convergence achieved using modest task sets. These improvements translate into superior individual parcellations and connectivity models. Together, our results dissociate reliability from validity in neuroimaging and challenge the prevailing assumption that rest provides a privileged window into intrinsic brain organization. Instead, functional architecture appears most faithfully revealed when the brain is actively driven through diverse task states.

## INTRODUCTION

Resting-state functional connectivity measured using *functional Magnetic Resonance Imaging* (fMRI) [1–11] has become one of the dominant methods used in human neuroscience. The approach relies on the fact that when participants lie in the MRI scanner, the *Blood-oxygen-level-dependent* (BOLD) signal spontaneously fluctuates around a mean (Figure 1a). From the co-variation of these fluctuations across regions, one can infer which regions are functionally connected. The approach can be used to infer the general organization of the human brain [9, 10, 12,13], but also to discover differences in brain organization across individuals [14–20]. Because resting-state imaging is easy to implement and can be obtained even from participants who cannot follow task instructions, it has become extremely popular in clinical and population neuroscience [2, 21–28]. However, one disadvantage is that the functional correlations between regions are driven not only by neural interactions, but also by noise sources, such as head movements, breathing, pulsatory heart-rate effects and scanning artifacts [29–34], which can bias correlation estimates.

**Figure 1.**
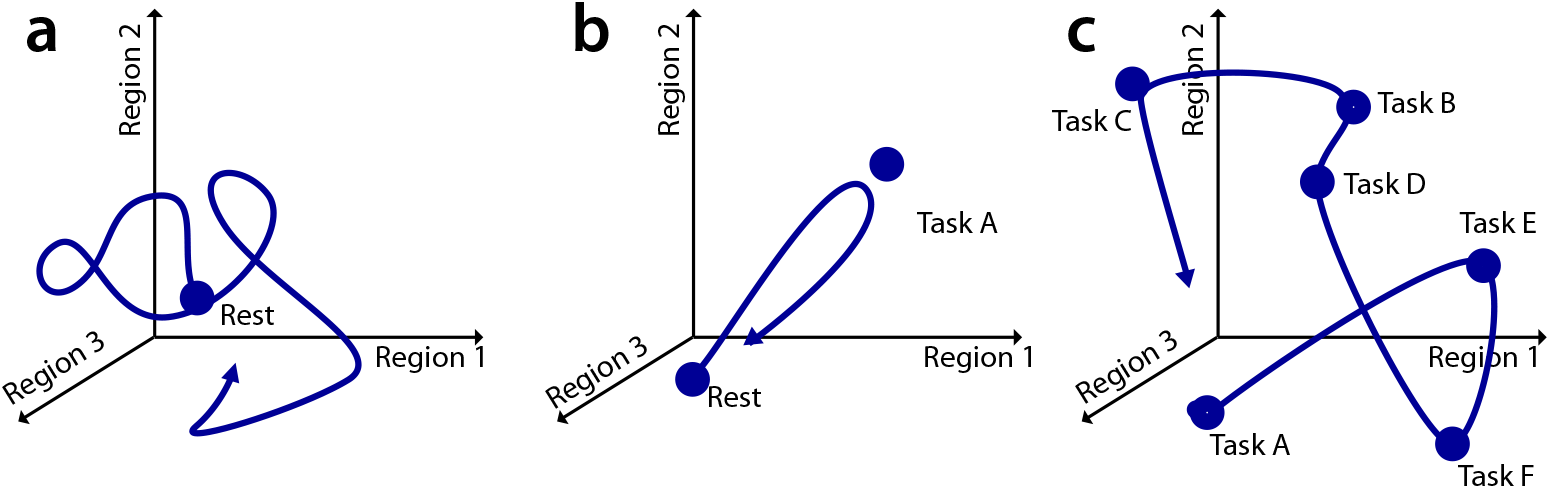
Hypothetical evolution of brain states, characterized by the activity in 3 brain regions. **(a)** Resting-state fMRI relies on spontaneous fluctuations around a mean point. **(b)** Task-based fMRI induces a replicable co-activation pattern across regions by alternating between a task and rest. **(c)** Multi-task fMRI drives the brain through a defined set of task-states in a random order.

In task-based fMRI (Figure 1b), participants typically alternate between short periods of rest and one or more task conditions. By using a randomized design, the experimenter can make strong causal inferences about how much of the variation in the measured BOLD activation is caused by the task, thereby retaining strong control of the measurement noise. However, the functional connectivity inferred from the task fMRI data will strongly depend on the co-activation patterns across the chosen tasks. For example, Task A may induce a strong co-activation (and hence correlation) between regions 1 and 2, whereas a different task may lead to a strong co-activation of 1 and 3 [35, 36]. Because of this task dependence, some researchers avoid task-based fMRI for connectivity studies, arguing that imposing specific cognitive or sensory states can bias the observed connectivity patterns and obscure the intrinsic functional organization of the brain [37–40]. When using task-based data to estimate functional connectivity, researchers often remove reliably induced task activation from task time series data and estimate functional correlations from the residuals [37, 41–46].

Multi-task fMRI (Figure 1c) provides a way to potentially avoid the disadvantages of both resting-state and task-based imaging. In this approach, brain activity is measured while participants perform a series of various tasks [20, 47–49]. If the sample of tasks is broad and representative enough, correlation estimates should become independent of the exact tasks chosen and converge toward an average task-independent functional connectivity. At the same time, the randomized design will provide strong control over the influence of measurement noise.

Here, we compared the estimated brain organization for individual participants from multi-task and resting-state fMRI studies. We evaluated these estimates by their ability to predict the individual’s brain organization across multiple novel tasks that were not included in the training data, since a good model of an individual’s functional brain organization should generalize across a wide variety of brain states. This choice reflects the ecological reality that the human brain predominantly occupies various *active* states in daily life, of which rest is only one.

There are many different statistical approaches to describe individual brain organization, including brain parcellations [10, 11, 50, 51], functional gradient mapping [52–56], network analysis [39,57 –61], and activity flow models [59, 62]. As evaluating all of these approaches in detail would be prohibitive, we chose the spatial (voxel-by-voxel or vertex-by-vertex) covariance matrix as our primary evaluation target. The spatial covariance matrix is a common point in the analysis of most brain organization models (Figure 2) and provides a summary of the raw data that contains all the necessary information to estimate most parcellations, functional gradients, and network models (i.e. it constitutes a sufficient statistic [63] for these models).

**Figure 2.**
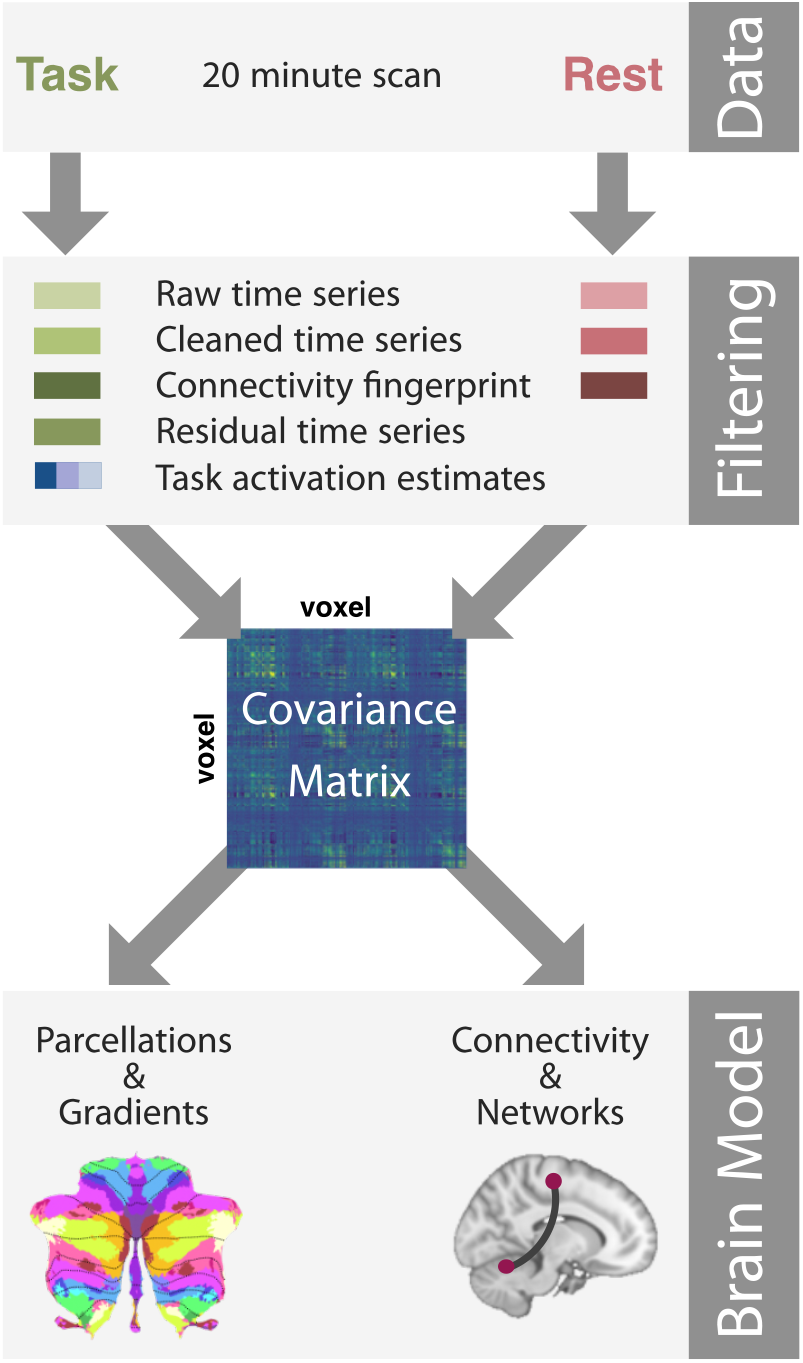
Overview of data analysis pipeline. **Data:** We used 20 minutes of task-based or resting-state fMRI data acquired from the same individuals. **Filtering:** Data were either used without further processing (*raw time series*) or subjected to one or more filtering steps: ICA-based denoising (*cleaned time series*), and subsequent connectivity fingerprinting (*conn. fingerprint*). For task data only, we used the experimental design to estimate task activation using a GLM approach (*Task activation estimates*). We also used the residual time series from this analysis (*residual time series*). **Covariance:** Voxel-wise or region-wise covariance matrices were then estimated for each subject and data type from the preprocessed time series data, connectivity fingerprints, or task activation estimates. **Brain model:** The spatial covariance can then be used to estimate a brain parcellation, gradient, or connectivity model.

In this paper, we estimate spatial covariance matrices from either task-based or resting-state fMRI data and compare the effect of different filtering approaches designed to reduce the impact of measurement noise. For task-based imaging, we also compare using either only the activation estimates for each task, the residuals from the time series model, or the entire time series. Overall we find that covariance matrices from multi-task designs generally outperform those based on resting-state data in predicting activity patterns in new, unrelated tasks. Although reducing the task-based time series to activation estimates decreases the reliability of covariance matrices, the predictive power is not reduced (or even superior) to those derived from the full time series. We then show that the results obtained at the level of spatial covariance matrices also generalize to two selected subsequent neuroimaging models: parcellating cortical and subcortical regions [11, 47, 64], and mapping their connectivity [65]. Finally, we replicate our results in two other datasets that contain both resting-state and multi-task fMRI data from the same individuals.

## RESULTS

In our first analysis, we used 20 minutes of resting-state and task-based data each, acquired from the same seventeen subjects in the *Multi-Domain Task Battery* (MDTB) dataset [47]. The multi-task data comprised 17 tasks tapping into cognitive, motor, perceptual, and social functions. During the resting-state session, subjects were instructed to look at a fixation cross in the center of the screen.

All time series data were first minimally preprocessed (see methods). We then compared a range of approaches to filtering or processing these data, which are commonly used in the field (**Figure 2**). First, we compared the impact of cleaning the time series data using *Independent Component Analysis* (ICA) [66–69] with that of using minimally preprocessed time series data. Second, we optionally used connectivity fingerprinting [70, 71] as an additional processing step. In this approach, the time series of each voxel was first correlated with the times series measured in 32 functional networks (see methods). Then, this vector of correlations (i.e., the connectivity fingerprint) was used instead of the time series as input data for subsequent analysis. This processing step effectively up-weights aspects of the time series that are commonly found in other brain networks, while suppressing aspects that are unique to the region. For task data, we also applied a standard *General Linear Model* (GLM) approach to estimate the evoked activation for each task in each run. We then used the vector of task activation estimates instead of the full time series for connectivity estimation. This approach concentrates on the part of the observed data that is reliably induced by the different tasks, and removes all fluctuation around the mean evoked activation. As a complementary approach, we also used the residual time series, which retains these fluctuations, while removing the activation that is systematically related to the tasks.

Subsequently, a spatial covariance matrix was computed from these different data types. This matrix provides an ideal measure for comparing different approaches, as it is the single common point for different filtering pipelines and different model types (Figure 2). In the main paper, we report the results for the neocortical covariance matrix. Confirmatory results were found for the cerebellar and cortico-cerebellar covariance matrices, as reported in the supplementary material. Finally, we show that our results generalize, as expected, to subsequent frameworks for parcellation and connectivity models.

### Reliability of covariance matrices

We first examined split-half reliability of the covariance matrices, estimated from two halves of neocortical data (10 minutes of scan time each). Although all data types showed reasonable reliability (all *r >* 0.57, one-sample t-tests all *t*_16_ *>* 13.25, all *p <* 3.87 × 10^−9^), we observed significant differences between the data types (Figure 3). Covariance matrices calculated from minimally preprocessed time series (task or rest) had low reliability and did not differ significantly from each other (*t*_16_ = 0.62, *p* = 0.543). ICA-based cleaning significantly increased the reliability of the covariance matrices (task raw vs task clean: *t*_16_ = −5.29, *p* = 7.27 × 10^−5^, rest raw vs rest clean: *t*_16_ = −7.40, *p* = 1.51 × 10^−6^). Subsequent connectivity fingerprinting led to a further increase both for rest and task time series; however, this was only significant for rest data (rest clean vs rest connectivity: *t*_16_ = −4.47, *p* = 3.85 × 10^−4^, task clean vs task connectivity: *t*_16_ = −1.82, *p* = 0.088).

**Figure 3.**
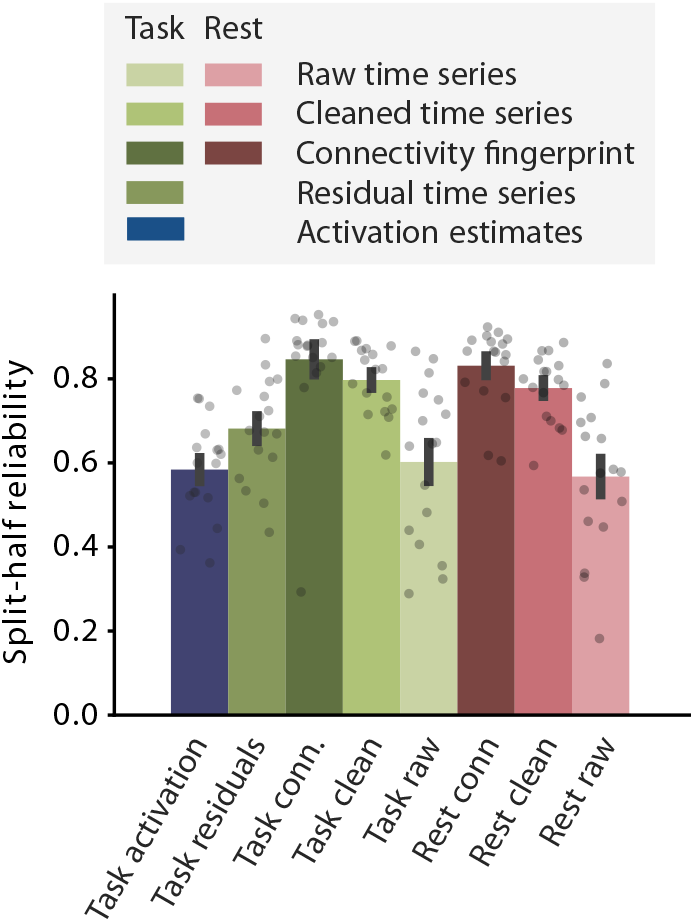
Split-half reliability of cortical covariance matrices. Split-half reliability was calculated as the correlation between the neocortical covariance matrices derived from 10 min of fMRI each. Covariance matrices are calculated from *Raw time series*: raw fMRI time series without further processing; *Cleaned time series*: fMRI time series after ICA-based denoising; *Conn. fingerprint*: Connectivity fingerprint calculated on cleaned time series data; *Residual time series*: Covariance matrix calculated on task residual time series (task-related activation regressed out); *Task activation estimates*: Activity estimates from a GLM for the 17 tasks. Error bars represent standard error of the mean across participants. Each dot is one participant.

In contrast, covariance matrices estimated only on the task-evoked activation had substantially lower split-half reliability than all data types but raw task and rest time series (all *t*_16_ *<* −3.12, *p <* 6.64 × 10^−3^; significant at a Bonferroni-adjusted threshold of (*p <* 0.007)). Even the residuals from the task-based time series showed significantly higher reliability. The split-half reliability of the cerebellar and cortico-cerebellar covariance matrices showed a similar pattern (Figure S1).

### Influence of measurement noise on covariance matrices

While reliability of covariance matrices is a necessary precondition to characterize stable functional organization in individual brains, it is not sufficient. In fact, high reliability of the covariance matrices can be due to measurement noise caused by head motion or physiological artifacts [29–32,72]. This is because noise processes can have consistent spatial structure: two voxels may be similarly influenced by a noise process, because their slice may be measured in the same relative phase of the cardiac cycle, or because they show similar brightness changes with residual head motion. These relationships are stable across multiple imaging runs, such that the correlation between the two voxels would be very reliable, even though it is caused by measurement noise.

In general, there is no guaranteed way to cleanly separate the covariance structure induced by measurement noise from that induced by neuronal processes. However, there are cases in which we can be relatively confident that high correlations must be induced by measurement noise. One example is the correlation between voxels in the superior cerebellum and voxels of the directly adjacent inferior occipital lobe. Even though these voxels are physically close, they are separated by the tentorium and not functionally coupled [73]. For this reason, previous analyses of cortico-cerebellar connectivity based on resting-state data forcefully removed these correlations [74]. We can therefore use the size of the correlation between voxels across the tentorium as a rough proxy of the degree to which different data types and processing pipelines induce artificially high correlations for spatially neighboring voxels.

We first confirmed that superior cerebellar voxels indeed show lower functional coupling with their immediate neighboring voxels in the inferior occipital lobe than with the rest of the neocortex by correlating the task-based activation estimates of cerebellar voxels within 10 mm of the tentorium with either the directly adjacent cortical voxels, or the remaining non-adjacent voxels (Figure 4a). To eliminate the influence of motion or physiologically induced noise correlations, we estimated the neocortical and cerebellar activation of different runs. A paired t-test of correlation differences showed that superior cerebellar voxels were less correlated with the immediately adjacent neocortical voxels than with the rest of the neocortex (mean r = -0.006 ± 0.004 vs. r = 0.014 ± 0.003, *t*_16_ = −4.52, *p* = 3.46 × 10^−4^). This confirms that the functional response of superior cerebellar voxels shows positive long-range correlations with the non-adjacent cortical regions, and significantly lower correlations with cortical regions that lie immediately across the tentorium.

**Figure 4.**
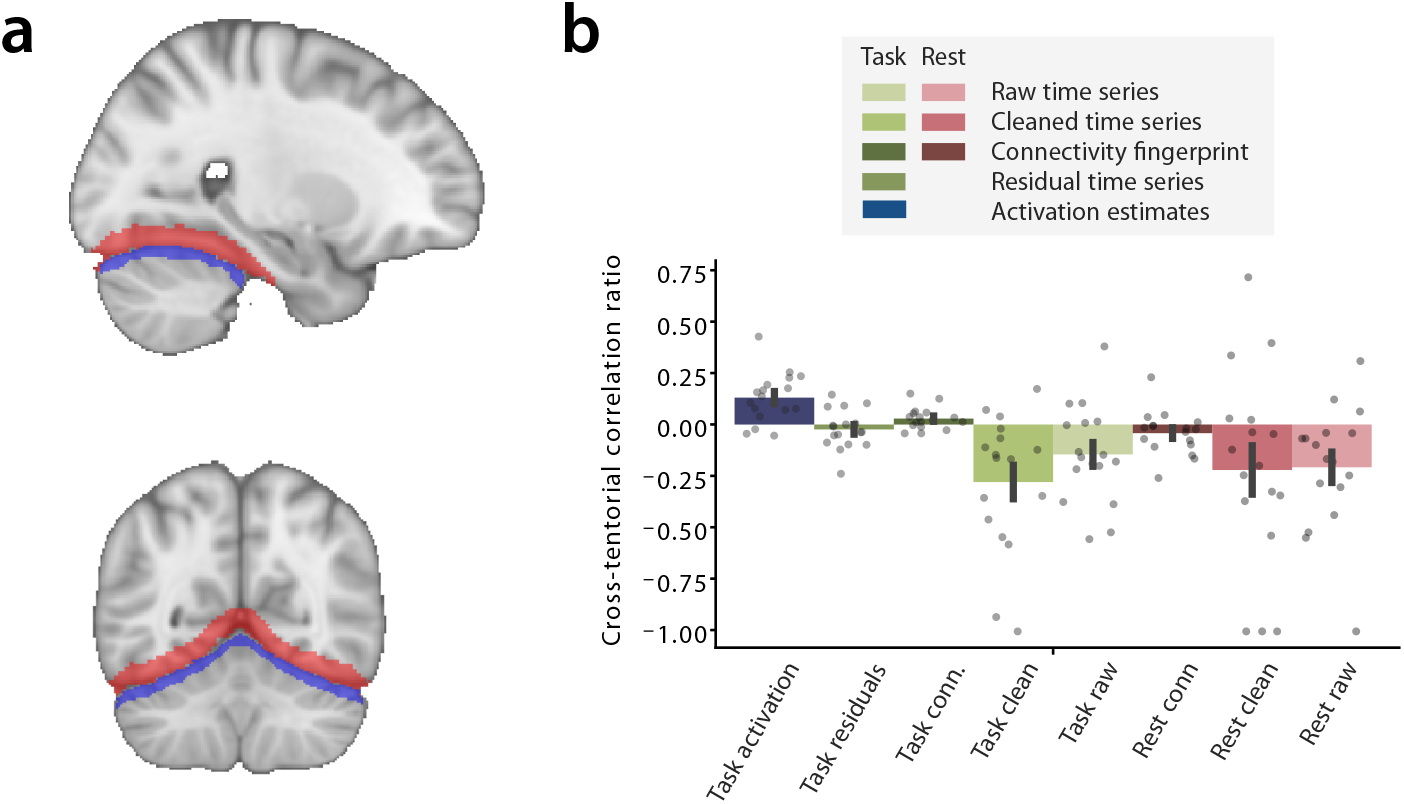
Normalized cross-tentorial correlation difference. **a**. Cross-tentorial correlations were calculated between superior cerebellar voxels (blue) and directly adjacent neocortical voxels (red, *r*_*a*_). Correlations were also calculated between superior cerebellar voxels and the rest of the neocortex (non-adjacent, *r*_*n*_). **b**. Cross-tentorial normalized correlation difference (*r*_*a*_ − *r*_*n*_)*/*(|*r*_*a*_ |− |*r*_*n*_|) for different data types. Error bars represent standard error of the mean across participants. Each dot represents one participant.

To examine how strongly different methods are influenced by shared noise, we calculated the normalized ratio between the correlations of the superior cerebellar and adjacent neocortical voxels and the superior cerebellar voxels and the rest of the neocortex (Figure 4a). This ratio should be positive for data that are less influenced by shared noise. We found that the correlation ratio for task activation estimates was significantly higher than zero (*t*_16_ = 4.42, *p* = 4.31 × 10^−4^, Figure 4b), and also exceeded the correlation ratio for all other data types (task activation estimates vs all other data types: all *t*_16_ *>* 2.99, *p <* 8.71 × 10^−3^). The correlation ratio for residual time series and connectivity fingerprints (calculated from task or resting-state data) did not differ significantly from zero (all *t*_16_ *<* 2.26, *p* = 0.305). The correlation ratios for raw and cleaned time series were negative, with the cleaned task time series showing a significantly lower correlation ratio than zero (*t*_16_ = −3.43, *p* = 3.41 × 10^−3^, Bonferroni-adjusted threshold of *p <* 0.007). This indicates that the time series of the superior cerebellar voxels is more correlated with the cortical voxels directly adjacent than with the rest of the brain. These correlations are likely driven by measurement noise that was not eliminated during preprocessing.

Hence, despite their relatively low split-half reliability, task activation estimates were least influenced by measurement noise that led to correlations of spatially proximal voxels. Raw and even cleaned time series from both task and rest data showed artificially inflated correlations between spatially adjacent voxels, a factor that will have contributed to their high reliability.

### Dependence of covariance matrices on the selected tasks

A common concern with using a task-based approach for deriving parcellations or connectivity models is that the results will be biased by the specific tasks used in the experiment. To quantify this effect, we examined the differences between the spatial covariance matrices derived from different task batteries, composed of 2, 3, 6, 10 and 13 tasks, sampled randomly from the 17 possible tasks. For each task battery, we used activation estimates that were based on 20 minute scan time each (or as close to 20 minutes as possible given the data available). The vector of resulting activation estimates at each brain location was then used to compute spatial covariance matrices.

We first examined the correlation of neocortical covariance matrices derived from task batteries consisting of 2 tasks. These were relatively different from each other, with correlations of 0.59 (± 0.12, Figure 5b). The systematic differences between the different task batteries can be visualized using classical *Multi-Dimensional Scaling* (MDS) (Figure 5a): each corner of the plot was occupied by task contrasts that emphasize the difference between different functional networks. Batteries contrasting rest with tasks associated with increased activation in the dorsal attention network and motor network (e.g., rest vs. N-back, rest vs. motor sequence) clustered in the lower right, whereas batteries involving action observation (e.g., action observation vs. motor sequence, action observation vs. N-back) clustered in the lower left quadrant. Batteries that contrast mentalizing against dorsal attention network (e.g., theory of mind vs. spatial navigation, theory of mind vs. N-back) in the upper left, while those contrasting scene construction and imagery (e.g., spatial navigation vs. GoNoGo, imagery vs. emotion recognition) occupied the upper right quadrant.

**Figure 5.**
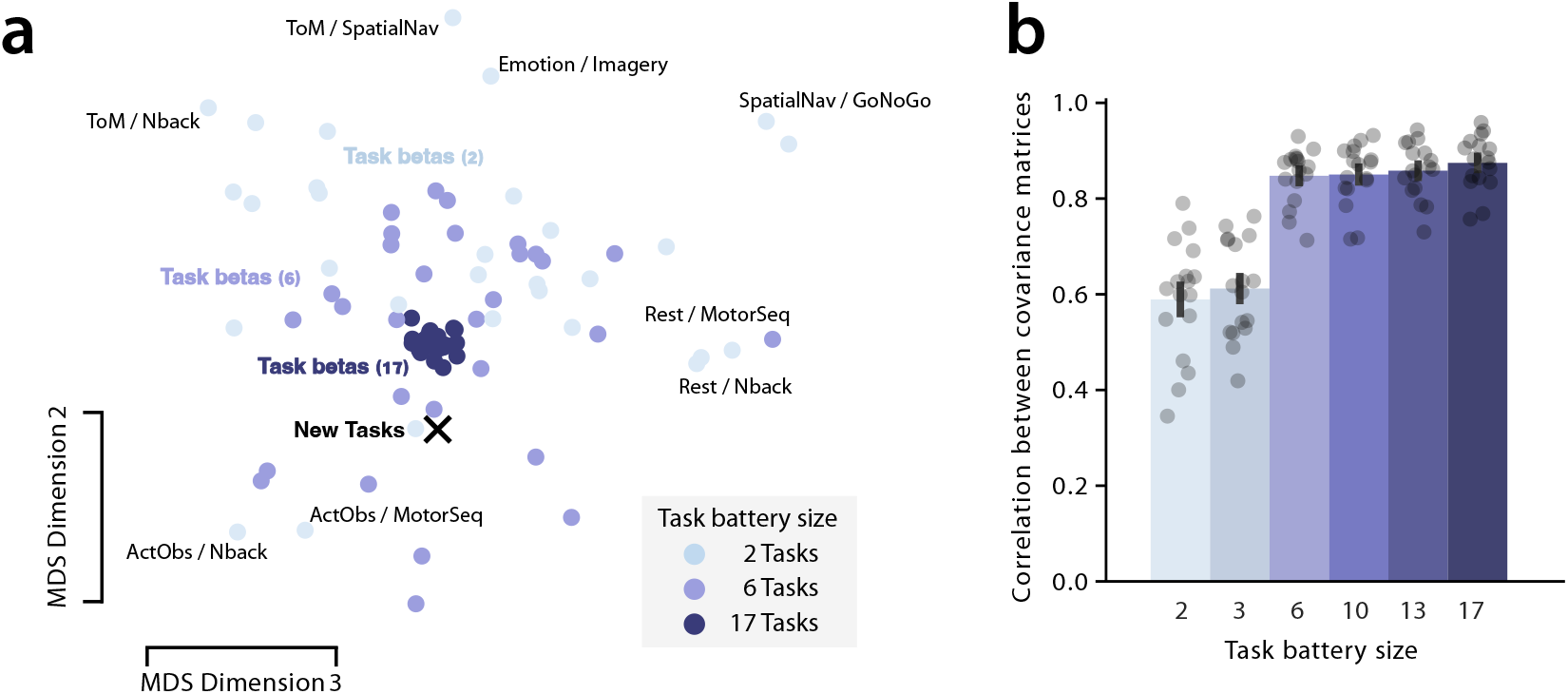
Similarity and differences between different task batteries. **a**. Multi-dimensional scaling graph of the similarity space of neocortical covariance matrices derived from different 20 min task batteries. **a**. The second and third MDS dimensions are displayed (for dimension 1, see Figure 6). Randomly sampled task batteries consisting of 2, 6 or 17 tasks are shown in progressively darker blue colours. Each dot shows the covariance matrix derived from a specific task battery averaged across subjects. All dots rely on independent data. The black X shows the covariance matrix derived from 18 novel task conditions, acquired during a separate session and without overlap to the sampled task batteries. Selected task batteries consisting of two tasks (a single task contrast) are labeled with abbreviated task names: *ActObs*: action observation task, *SpatNav*: spatial navigation task, *GoNoGo*: Go-Nogo task, *Imagery*: Motor imagery task, *ToM*: theory of mind task, *MotorSeq*: motor sequence task, *Nback*: N-back working memory task, *Emotion*: emotion recognition task. **b**. Average similarity of randomly sampled task batteries to each other. Each dot represents the within-subject, pairwise correlations between the covariance matrices.

However, the systematic difference between different task batteries could be significantly reduced by including more randomly sampled tasks in the battery. As we increased the number of tasks from 2 to 6, we observed a substantial increase in the correlation (Figure 5b). Increasing the number of tasks to 10, 13 and 17 tasks did not lead to substantial further improvements (17 tasks vs. 10 or 13 tasks: all *t*_16_ *<* −1.42, *p >* 0.17). All 17-task batteries contained the same set of tasks. Consequently, similarity at 17 tasks was constrained primarily by measurement reliability, as the correlated covariance matrices were estimated from independent data sets. In sum, these findings show that the more varied the task battery is, the more invariant the correlations between brain locations are to the exact choice of tasks, with batteries comprising six or more tasks approaching the noise ceiling.

### Generalization to novel task set

Critically, a good model of an individual’s functional brain organization should not only be reliable and invariant to the exact battery chosen but should also make valid predictions across a wide variety of brain states. We therefore evaluated how similar the covariance matrix derived from each different data type was to the covariance matrix obtained from the same individual, in a separate session comprising 18 new task conditions and 144 minutes of functional imaging data. By calculating the similarity as the Pearson correlation between the vectorized covariance matrices, we were able to situate the covariance matrix of the independent task set in the MDS space (Figure 5a, 6b).

The MDS plot showing dimension 2 and 3 (Figure 5a) clearly shows that the covariance matrix of the new task set falls quite close to the center of the randomly sampled task sets, confirming our claim that covariance matrices (and hence brain connectivity and parcellations models) estimated from rich task batteries converge on a common, invariant point per individual. Indeed, the average correlation between the new task set and the 20-minute 17-task batteries were highest with r= 0.56 ± 0.11, with slightly declining similarity when reducing the number of tasks. While there was no difference between 17 and 13 tasks that survived the Bonferroni-adjusted threshold of *p <* 0.01 (13 tasks: 0.56 ± 0.10, *t*_16_ = 2.90, *p* = 1.04 × 10^−2^), progressively reducing the number of sampled tasks led to significant decreases in similarity to the new task set (all *t*_16_ *<* −5.46, *p <* 5.20 × 10^−5^).

So far we have only compared covariance matrices estimated on tasks-activation data, with differences relating mostly to the exact choice of tasks. These differences were well represented in the second and third dimension of the MDS space. In contrast, the first dimension of the MDS space (Figure 6a) differentiates between covariance matrices based on task activation estimates and those estimated from the entire time series.

**Figure 6.**
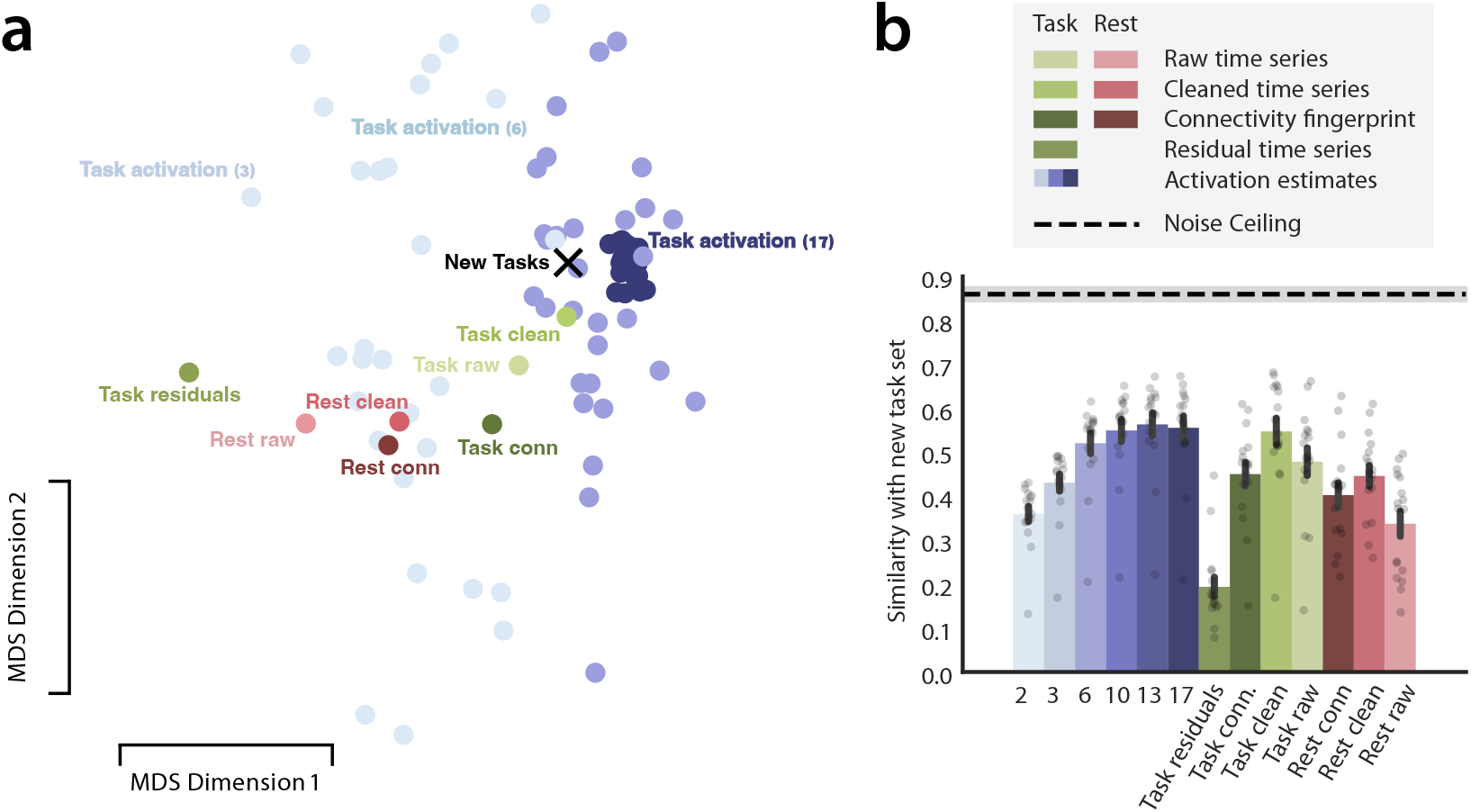
Similarity space of covariance matrices. **a**. First two dimensions of the similarity space of neocortical covariance matrices. *Rest raw*: raw resting-state time series, *Rest clean*: cleaned resting-state time series, *Task raw*: raw task time series, *Task clean*: cleaned task time series, *Task residuals*: residual task time series, *Task conn*: connectivity fingerprint calculated on task time series, *Rest conn*: connectivity fingerprint calculated on resting-state time series, *New Tasks*: task activation estimates estimated from a separate session comprising 18 unique task conditions. Scan time of all data types was matched to 20 minutes, except for the independent new task session, which was 144 minutes per individual. Similarities were calculated within participants then averaged across participants. For task batteries, each dot represents one specific sampled task battery. **b**. Correlations of covariance matrices with the same individual’s new task session. Noise ceiling was estimated by computing the similarity between covariance matrices derived from two runs and covariance matrices derived from the remaining fourteen runs of the independent task session on which all similarities were evaluated. Error bars represent standard error of the mean across participants. Each dot represent one participant.

On the left side of this axis was the covariance matrix estimated from raw resting-state time series, which had the lowest similarity with the independent task session (r=0.34 ± 0.11). ICA-based cleaning resulted in an estimate that was significantly closer (r=0.45 ± 0.10, Figure 6b, *t*_16_ = 6.08, *p* = 1.60 × 10^−5^). On the other hand, connectivity fingerprinting did not improve, but decreased the similarity to the covariance matrix estimated on an independent task set (rest clean vs. rest connectivity: *t*_16_ = −4.29, *p* = 5.64 × 10^−4^).

Importantly, just by using time series data that were collected while the individual was engaged in a variety of tasks (but otherwise processed like resting-state data), we obtained an estimate that was more predictive of the covariances during a completely different set of tasks. The raw and cleaned task time series data were significantly more similar to the independent task session than the corresponding estimates from the resting-state data (raw task vs. raw rest: *t*_16_ = 3.60, *p* = 7.20 × 10^−3^, cleaned task vs cleaned rest: *t*_16_ = 2.95, *p* = 2.84 × 10^−2^). These findings show that the activation caused by the broad set of tasks allows for better identification of a task-invariant connectivity pattern than using data acquired during rest, no matter how the corresponding time series was processed.

Surprisingly, the task-based covariance matrices were just as predictive when calculated on task activation estimates only (*t*_16_ = 0.57, *p* = 0.57), even though this step removes a lot of information from the data and reduces the reliability of the covariance matrices (Figure 3). Conversely, using only the information that has been removed, i.e. the residuals from the regression, led to a covariance matrix that was significantly less similar to the new task set than the covariance matrix estimated on the entire time series (*t*_16_ = −9.52, *p* = 5.46 × 10^−8^). These results indicate that the predictive information that generalizes well to new task states is mostly captured in the average evoked activation and to a much lesser extent in the fluctuations around this mean.

For cerebellum and cortico-cerebellar covariance matrices, a similar pattern emerged (supplementary results, Figure S3). Furthermore, to ensure that results are not restricted to the use of a spatial covariance matrix as the main evaluation target, we showed they generalized to the spatial correlation matrix (Fig. S4). Finally, we also replicated our analysis using the task and rest data from the *Human Connectome Project* [75, 76] *(HCP, Figure S5)*.

In sum, these findings show that multi-task data consistently captured functional organization better than rest data, irrespective of preprocessing. Most of this useful information was captured in the mean evoked activation. The advantage of the multi-task approach was independent of matrix statistic (covariance or correlation), brain structure (cortex or cerebellum), and dataset.

### Application to individual brain parcellation models

High similarity to the covariance matrix estimated on a new task set suggests that any model estimated on this sufficient statistic will be able to better predict this new task data. To demonstrate this intuition directly, we compared the performance of different data types on two common neuroimaging problems: deriving individual functional parcellations [20, 51, 77], and estimating models of connectivity [65].

For the parcellations, we evaluated how well individual maps—derived from each data type—predicted the functional boundaries measured in an independent set of tasks. We performed this analysis in two brain structures that tend to show a high inter-individual variability in organization: the prefrontal cortex and the cerebellar cortex [78]. Parcellations were derived using a previously published probabilistic parcellation algorithm (see Methods,[77]). As priors, we used 44 regions of the Glasser parcellation [11, 79] in each hemisphere for the prefrontal cortex and the 32 functional region of a recent parcellations of the cerebellar cortex [64]. For each subject, we evaluated how well the winner-take-all map predicted functional boundaries in held-out data from the same subject using the *Distance-Controlled Boundary Coefficient* (DCBC, [80] - see Methods - DCBC evaluation).

In the prefrontal cortex, an individual parcellation based on the task activation estimates achieved the highest DCBC (mean DCBC = 0.18 ± 0.05), numerically outperforming all other data types (Figure 7a). Task activation estimates resulted in significantly higher DCBC than all data types (all *t*_16_ = 4.08, *p* = 8.79 × 10^−4^), apart from the task connectivity fingerprint (*t*_16_ = 0.68, *p* = 0.504). As predicted by the similarity of covariance matrices, task data again outperformed rest data for matched preprocessing (all *t*_16_ *>* 2.77, *p <* 1.36 × 10^−2^).

**Figure 7.**
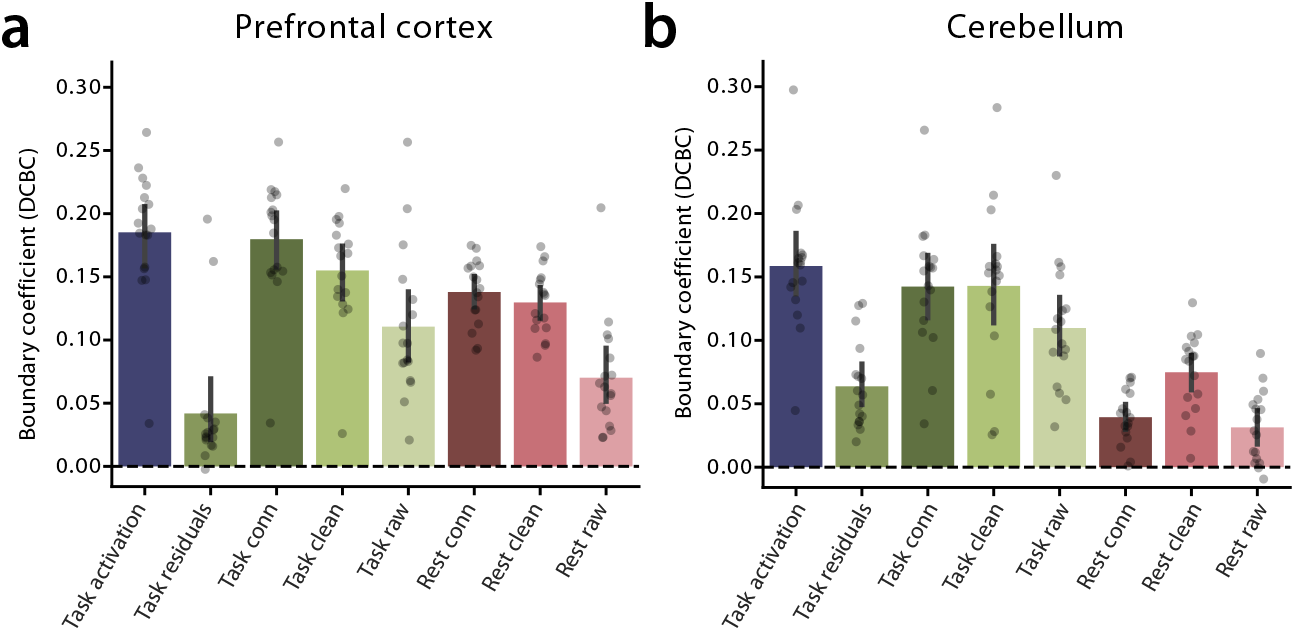
Performance of individual parcellation maps in predicting functional boundaries in independent tasks for the same subjects in **a**. Prefrontal cortex, **b**. Cerebellar cortex. The Distance controlled boundary coefficient (DCBC) indicates the difference of the within-parcel vs. between-parcel correlation, controlled for the spatial distance of two brain locations. For random, but spatially contiguous parcellations, the expected DCBC is zero.

In the cerebellum, the results closely mirrored the prefrontal cortex: task activation estimates parcellations yielded the highest DCBC (mean DCBC = 0.16 ± 0.05), and significantly outperformed all other data types (all *t*_16_ *>* 3.21, *p <* 5.45 × 10^−3^), apart from the cleaned task timeseries (*t*_16_ = 1.58, *p* = 0.13). As expected, task data outperformed rest data for matched preprocessing (all *t*_16_ *>* 4.77, *p <* 2.08 × 10^−4^). For full results, see Supplementary Materials section S7.

In addition to our main multi-task data set, we repeated the analysis on a separate replication dataset (N=8, see Methods). This dataset contained 40 minutes of multi-task fMRI and 40 minutes of rest fMRI to parcellate the cerebellum, and 40 minutes more of multi-task data to evaluate the functional boundaries. We found the same pattern of results (Figure S6), with parcellations derived from task activation estimates showing highest performance (all *t*_7_ *>* 3.56, *p <* 9.20 × 10^−3^). Task data resulted in significantly higher performance than rest data for matched preprocessing (all *t*_16_ *>* 3.31, *p <* 1.30 × 10^−2^). For full results, see Supplementary Materials section S8.

Together, these results demonstrate that parcellations estimated from multi-task data, especially when using only task activation estimates, provided the most accurate description of individual functional boundaries in both prefrontal cortex and cerebellum, as these predicted the boundaries measured under novel task states the best. As before, data from multi-task fMRI were more powerful than the data from resting-state scans, even if the data were analyzed exactly the same way in both cases (i.e. treated like resting-state data). While cleaning and fingerprinting could improve the prediction of individual functional boundaries, it never exceeded the level achieved when using activation estimates only. This pattern of results was independent of brain structure and replicated in a separate dataset.

### Application to individual brain connectivity models

As an example for a connectivity model, we chose to model the connectivity between two highly interconnected brain areas, the neocortex and the cerebellum [65]. We estimated a directed partial connectivity model that used the entire pattern of the cortical fMRI activation to predict the activation in each cerebellar voxel separately. We trained individual linear task-invariant regression models [65, 81] on each data type and evaluated how well the model can predict the cerebellar activation in the same individual in a new set of tasks (see Methods - connectivity modeling).

In predicting the cerebellar activation pattern on novel tasks, connectivity models based on task activation estimates achieved the highest predictive correlation (mean *r* = 0.31 *±* 0.04, Figure 8), significantly outperforming other data types (all *t*_16_ *>* 2.70, *p <* 7.82 × 10^−3^, Bonferroni-adjusted threshold *p <* 0.01). Also, task data outperformed rest data for matched preprocessing (all *t*_16_ *>* 7.82, *p <* 3.70 × 10^−7^). Finally, cleaning increased performance of the task and rest over raw time series.

**Figure 8.**
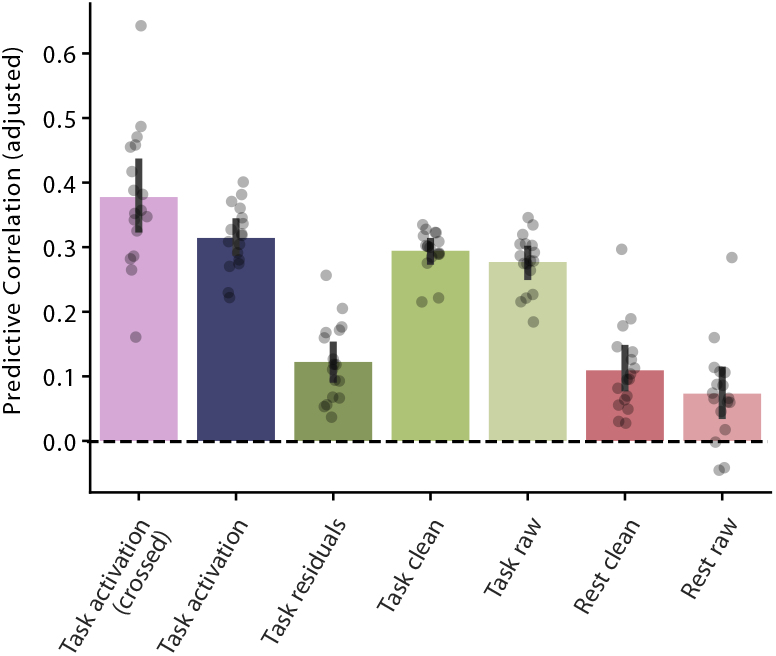
Connectivity model results. Shown is the correlation of predicted and observed cerebellar activation patterns. Predictions are based on the individual connectivity models (trained for each data type) and the individual neocortical activation patterns for a new set of tasks. Each dot represents a single subject.

In the original paper [65], we employed a *crossed* approach to reduce the influence of measurement noise onto the estimated connectivity weights, i.e. we predicted cerebellar activation patterns from one imaging run using cortical activation patterns from the same tasks acquired in a separate imaging run. We found that training on one run and testing on the other further enhanced the predictive performance: the crossed model (mean *r* = 0.38 *±* 0.03) significantly outperformed the model based on uncrossed task activation estimates (*t*_16_ = 2.34, *p* = 1.64 × 10^−2^). This suggests that even connectivity models based on task activation maps are biased to some degree by measurement noise. Estimating these in cross-validated fashion reduces this biasing influence and improves predictive performance for novel sets of tasks.

## DISCUSSION

### Summary

In this study, we systematically compared resting-state and multi-task approaches to capture individual functional brain organization. We demonstrated that the spatial covariance and correlation matrices based on multi-task data were more similar than those estimated on resting-state data to the covariance or correlation matrices measured under novel task conditions. This was even the case when the time series data for rest- and multi-task data was pre-processed and analyzed exactly in the same way. As expected, ICA-based cleaning [67–69] and sometimes also connectivity fingerprinting [70,71], improved the similarity of these connectivity matrices further. Surprisingly, the advantageous features of the covariance matrices estimated from task data were fully preserved, or even improved, when we only used the averaged task-evoked activation maps, rather than the entire time series.

Our results also show that the advantage of multi-task fMRI is only present when the battery contains enough tasks. Task-based data relying on a few tasks bias the resulting activation strongly towards the task set and do not generalize well to novel tasks. However, 6 or more diverse tasks appeared to be sufficient to obtain a reasonable estimate of the task-invariant correlation structure of the brain. Our findings on covariance matrices directly translated to models that were based on these matrices: multi-task data outperformed rest data for individual parcellations and connectivity modeling, with task activation estimates showing the highest performance. Finally, we replicated these findings in an independent high-resolution dataset, in the HCP task-dataset, and across various brain structures, confirming that our results are not driven by the idiosyncrasies of a single dataset.

### Reliability does not guarantee good predictive performance

Our study also demonstrates that the reliability of connectivity matrices is a necessary, but not sufficient, condition for good predictive performance. Individual connectivity matrices calculated on cleaned time series data or on connectivity fingerprints tend to show relatively high split-half reliability [16, 18, 45, 82– 84], a result that we replicate here. In contrast, connectivity matrices based on task activation estimates alone were much less reliable, likely resulting from the fact that a large amount of the data are discarded when projecting the time series onto a design matrix to obtain activation estimates. However, task activation estimates best generalize to novel tasks, suggesting that they provide a more valid estimate of task-invariant connectivity. In contrast, the high reliability of spatial covariance matrices estimated from time series data does not guarantee that the resulting connectivity analyses or brain parcellations are valid.

We suggest that the high reliability of time series data is, at least to some extent, caused by spatially correlated physiological or movement-induced measurement noise. While the signals induced by the noise processes are themselves uncorrelated across runs, spatial-correlation matrices calculated based on them are not, but can show up as very stable signal correlations across independent measurement runs.

Our cross-tentorial analysis provides some evidence for this. Task activation estimates in the superior cerebellum were less correlated with immediately adjacent neocortical voxels than with functionally connected, but spatially distant regions. In contrast, raw and cleaned time series from both task and rest data showed higher correlations with immediately adjacent neocortical voxels as compared to the rest of the brain. Because there are no known functional connections between the cerebellum and the immediately abutting neocortex—and the two are separated by a hard membrane—it is likely that these correlations were induced by shared measurement noise. In previous cortical-cerebellar connectivity models, authors have therefore chosen to forcefully remove these correlations by regressing out the signals from the abutting neocortex in the cerebellum [74]. However, the shared noise correlations across the cerebellar tentorium are very likely also present elsewhere in the human brain, where they, unfortunately, can not be distinguished from real neuronal correlations between adjacent brain regions. Reducing the multi-task time series to activation estimates appears to remove these correlations effectively.

### Task diversity reduces bias

A common concern with task-based approaches is that the resulting brain organization will be biased toward the specific tasks used [37, 41–46]. Our results clearly demonstrate that this concern is fully warranted: Depending on the type of task contrast, covariance matrices estimated on only 2 tasks differed strongly and systematically from each other, as visualized in their location in the MDS similarity space.

However, we also demonstrate that this bias can be effectively reduced when including more tasks, even if the scan time is held constant, such that we have much less data per task. The similarity between different task batteries increased significantly when going from 2 to 6 tasks, but then started to plateau, with no significant improvement beyond 6 tasks. This result may partly reflect the limitation that we could only sample from 17 different tasks, such that larger task batteries include overlapping sets of tasks. However, we also showed that the generalization of the results to a completely novel task set plateaued after including 13 tasks, and that relatively good performance could already be reached with 6-10 tasks. This suggests that a good estimate of a task-invariant individual parcellation or connectivity model can be achieved using relatively small task batteries (6–10 tasks).

### Active multi-task engagement helps to reveal task-invariant functional organization

Multi-task data consistently outperformed rest data across all matched preprocessing steps, from raw time series to cleaned time series to connectivity fingerprints. This robust result suggests that the advantage of using a multi-task battery does not lie in how the information is used in subsequent analyses, but in the information content of rest versus task data. Active engagement in diverse tasks drives the brain through a series of states, which seems to provide richer, more informative signals about functional organization than the spontaneous fluctuations recorded during rest, regardless of how the data are processed.

### Using activation estimates or residuals from the task-based time series

The full task time series can be decomposed into two complementary aspects: The mean evoked activation maps for each task (the task activation estimates), and the random fluctuations around this mean (the residual time series). Previous work has argued strongly that for building connectivity models from task data, the task-related activation should be removed and only the residual time series be used [37, 41– 46]. This step is aimed at reducing the dependency of the result of the specific choice of tasks. It has been further shown that regressing task activation out yields maps that resemble resting-state-derived connectivity [37] and that can recover within-individual network architectures close to those estimated from resting-state [85]. We replicate these findings conceptually, by showing that covariance matrices from residual time series are highly reliable and can approximate individual functional organization at a level similar to resting-state time series. Furthermore, on all our evaluation criteria (similarity to novel tasks, parcellation, connectivity), the performance of residual-based estimates exceed zero, which indicates they indeed carry relevant signal about the individual.

However, we also find that despite good reliability and similarity to rest, the residual time time series performed substantially worse than both the full task time series and the task activation estimates alone in predicting functional organization in unseen tasks. It is likely that this behavior is due to the biasing effect of measurement noise: Our cross-tentorial analysis indeed suggests that correlation estimates based on residual time series, despite being highly reliable, are more susceptible to shared noise. This pattern is consistent with the view that residuals reflect a mix of useful fluctuation and measurement noise. While the residual time series contain some signal about individual functional organization, these signals are diluted, even after cleaning, with measurement noise that biases the estimated covariance structure. Importantly, Gratton et al. [45] demonstrated that functional networks are dominated by stable group and individual factors rather than cognitive state; our results extend this by showing that when the goal is *prediction* of organization during novel task states, the task-evoked activation from a diverse battery captures that stable organization more validly than do rest or residual-based estimates.

Overall, therefore, when applied in the context of a rich multi-task battery, regressing out task-related activation substantially reduces the similarity of the spatial covariance matrix to other task sets. This suggests that the information that reveals the task-invariant organization of an individual brain primarily resides in the task-evoked responses themselves, rather than in the residual spontaneous fluctuations.

### Theoretical implications for functional precision mapping

Our findings challenge the assumption that BOLD fluctuations observed during rest reveal the “true” or ‘baseline” functional organization. Instead, functional organization may be better characterized when the system is actively driven through diverse states. Diverse task states seem to provide an efficient way to sample the brain’s intrinsic functional organization, yielding representations that generalize to other functional contexts. This aligns with the idea that the brain’s functional architecture is optimized for action, not rest.

Recent reviews of resting-state fMRI have emphasized both the transformative impact of rest data on network neuroscience and the limitations that arise when reproducibility and reliability are treated as proxies for functional relevance [2–4, 13, 39, 86]. Although resting-state data can robustly capture some intrinsic functional organization [8, 45, 87, 88], and Smith et al. showed close correspondence between the brain’s functional architecture during activation and rest [8, 89], accumulating evidence suggests that rest data is not optimal for all neuroscientific questions, particularly those involving prediction of behaviour [20, 90–93] or generalization to novel cognitive states [94]. Our findings provide a concrete empirical example of this broader point: although resting-state data are highly reliable, they are less effective than task-based estimates at predicting functional organization during novel cognitive states. Gratton et al. [45] showed that functional networks are dominated by stable individual factors rather than cognitive state; here we show that for *predicting* that organization in new task states, multi-task activation estimates outperform both rest and task-residual estimates. Consistent with longstanding concerns about spurious correlations driven by spatial proximity and physiological confounds in rest data [31, 32], we show that even state-of-the-art cleaning approaches do not completely remove the biasing effect of structured noise, whereas using task-evoked activation estimates substantially reduce these effects. These findings support a growing view that functional organization should be evaluated not only by stability, but by its capacity to generalize to functional organization in other contexts. Overall, therefore, multi-task fMRI provides a powerful alternative to resting-state approaches for characterizing behaviourally relevant brain organization.

Our findings also highlight the importance of choosing appropriate validation metrics. While reliability is necessary, it is not sufficient. Internal validity metrics, such as the cross-tentorial correlation ratio, provide crucial sanity checks that reliability alone cannot provide. When the goal is prediction or generalization, validity metrics should be prioritized alongside reliability metrics. The present results align with recent concerns about generalization failure in rest-based analyses: representations that are stable and reproducible do not necessarily support extrapolation to new cognitive states [2, 94–96]. Evaluating functional organization by its ability to predict unseen tasks may therefore provide a more stringent and functionally meaningful benchmark.

### Practical implications for functional precision mapping

If the goal is to map the intrinsic functional brain organization of individuals, we recommend using a diverse task battery instead of acquiring resting-state data. In order to provide a general mapping that generalizes well to other tasks, our data show that such a battery needs to sample different task states broadly. We found that the spatial covariance matrix stabilized with task batteries that contained more than 6 diverse tasks. Thus, even if functional scan time per individual is restricted, as is often the case in clinical or population-based studies, it is possible to acquire multiple repetitions of a battery when switching between tasks every 35s [47]. For example, within 15 min of functional scan time, it is possible to acquire 3 independent runs with a battery size of 8 tasks. While the exact choice of tasks will often depend on practical considerations, it is imperative to choose an efficient task set that is able to drive the brain through as many different states as possible. We offer a quantitative approach to optimal battery design, along with practical recommendations for implementation, in a companion paper [97]. That paper also describes a Python library with many pre-implemented tasks, which makes it easy to design and run such batteries.

Once the data is acquired, it can be used in two ways: First, the time series acquired during task fMRI can be used in the same way as a resting-state time series to estimate intrinsic network organization and connectivity structure [20, 58, 64, 65, 77, 85]. In contrast to using task data with a single task contrast [37, 41, 45], the task-evoked component in a multi-task fMRI design does not have to be removed. Rather we show that using the entire time series yields superior connectivity models compared to resting-state or residual data alone.

Additionally, the data can be analyzed with a GLM approach to estimate task activation maps for each run. The activation profile across these tasks provides an independent way of characterizing the functional networks. In resting-state data, the correspondence of networks across individuals has to be made by the spatial location of the networks [45, 98]. The task-activation profiles now anchor the analysis in an interpretable way and allow for strong comparison between individuals, allowing researchers to clearly identify and separate even networks that are spatially highly interdigitated [99, 100]. Furthermore, by calculating the reliability of task-evoked activation maps across runs, the researcher has an independent assessment of the reliability of the data, which is not inflated by spatially structured noise.

Of course, in some contexts, resting-state approaches still hold great value for mapping intrinsic architecture. It is uniquely practical for infants, clinical groups and other populations that cannot perform tasks, and provides an important tool for cross-species comparisons [2, 21–28,101,102]. Still, when the goal is to capture behaviorally relevant, generalizable organization, multi-task fMRI appears superior to rest data alone.

### Conclusion

We demonstrate that multi-task fMRI provides better predictions of the task-invariant functional brain organization than resting-state fMRI. This advantage persists across preprocessing steps and translates directly to common neuroimaging applications like parcellation and connectivity modeling. While multi-task fMRI can be challenging when measuring many different tasks, our findings show that task batteries of 6 or more tasks already start to converge towards a good task-invariant estimate, making multi-task approaches feasible. These results challenge the assumption that rest provides a privileged window into task-invariant brain organization. Instead we suggest that the functional organization of the brain is best assessed by actively driving it through a series of well-controlled and diverse multi-task states.

## METHODS

### Participants

We used two locally acquired datasets that included multi-task and resting-state fMRI data from the same subjects for the main analysis. The task sessions of each dataset comprised a broad battery of tasks tapping into cognitive, motor, perceptual, and social functions: (1) The *Multi-Domain Task Battery* dataset (MDTB, N = 17 [47]), (2) a high-resolution *Replication dataset* (N = 8 [97]).

All participants gave informed consent under an experimental protocol approved by the institutional review board at Western University. None of the participants reported a history of neurological or psychiatric disorders or current use of psychoactive medications. The MDTB data set had 10 female and 7 male individuals (mean age = 24, sem=0.71); the Replication data set had 4 female and 4 male individuals (mean age= 22.63, sem = 0.57).

Finally, we used the task and rest data from the *Human Connectome Project* (HCP) Unrelated 100 subjects which is publicly available in the HCP S1200 2017 release [75].

### Data acquisition

#### MDTB Dataset

Acquisition parameters for the MDTB dataset have been reported in [47]. In brief, twenty-four participants (16 female, mean age = 23.8) were scanned using a 3T Siemens Prisma scanner at the Center for Functional and Metabolic Mapping at Western University. Task-based fMRI data was acquired during 4 task sessions using a multi-band EPI sequence (TR = 1s, voxel resolution = 2.5 ! 2.5 ! 3.0 mm, multi-band acceleration factor 2, in-plane acceleration factor 3, 598 volumes per run). Resting-state fMRI data was recorded using the same scanning parameters for seventeen of the participants while they fixated on a white crosshair on a grey background for 2 runs. In this paper, we only consider data from the seventeen participants with both task and rest fMRI data.

We conducted five scanning sessions, separated at least by a day. In the first four scanning sessions, participants performed eight task-based runs each. The first two sessions were conducted using task set A, and the second two sessions were conducted with task set B. Each task set contained 17 tasks spanning motor, cognitive, language and social domains. Each task block included a 5-second instruction period and 30 seconds of performing the task. This resulted in 16 imaging runs of 10 minutes for each task set. In the fifth session we acquired two 10 min runs of resting-state data.

In this paper we used task set A (or the resting-state scans) as a training set to estimate parcellation or connectivity models, and task set B as our test set to assess how well these models generalized to a new set of tasks. We excluded eight tasks (including rest) from the test data, since they were also presented in task set A. This ensured that we could test the generalization of the models to a fully new set of tasks. A detailed description of the tasks in each task set after exclusion can be found in Supplementary Table 1 and 2.

#### Replication Dataset

Participants were scanned using a 7T Siemens MAGNETOM MRI Plus scanner at the Center for Functional and Metabolic Mapping at Western University. Eight runs of task-based fMRI data was acquired during one task session using a multi-band EPI sequence (TR = 1.1s, voxel resolution = 2.3 mm isotropic, 523 volumes per run, multi-band acceleration factor 2, in-plane acceleration factor 3). Seven or eight runs of resting-state fMRI data of the same length were recorded using the same scanning parameters while they fixated on a white crosshair on a grey background. Task and rest scanning acquisition took place on a different day. A structural T1-weighted MRI scan was acquired at the beginning of the task session using a magnetisation prepared rapid acquisition gradient echo (MPRAGE) sequence (TR = 2200 ms, voxel resolution = 0.7 mm isotropic). For four subjects, the scanning parameters of the task session differed slightly from the rest (TR = 1.3s, volumes = 443).

During the task session, participants performed 16 tasks during eight 10-minute imaging runs. Each task was performed once during each run, with task order randomized across subjects and runs. Within each run, after a 5-second instruction period, participants performed the task for 30 seconds. For a detailed description of the tasks, see Supplementary table 3 [97].

#### HCP Dataset

Data acquisition parameters for the HCP datasets can be found in Van Essen et al. [75]. We used the minimally preprocessed 3T task-fMRI data from the HCP 1200 subjects 2017 release [75], selecting 50 of the 100 unrelated-subjects group (25 female, 25 male, mean age estimated based on age ranges = 29.7). The task data included two imaging runs for each of seven task sessions (working memory, gambling, motor,language, social, relational and emotion), where each of the imaging runs was 2.5-5 minutes long. The rest data included four imaging runs of 15 minutes each.

### Functional Data preprocessing

Both task-based and resting-state functional data was preprocessed as described in Zhi et al. [77]. Preprocessing steps included correction for spatial distortions, motion correction and spatial coregistration to the individual anatomical [103]. No smoothing or group normalization was applied at this stage. For the HCP resting-state data, cleaning was already applied by default. Using a unified code framework (available at github.com/diedrichsenlab/Functional_Fusion), the data were then extracted in two atlas spaces. For the neocortex, we reconstructed the individual white-matter-to-gray-matter surface, as well as the pial surface using the recon-all pipeline of the freesurfer package [104, 105]. These individual surfaces were aligned and resampled to the symmetric freesurfer32k template [106]. Functional data were projected onto the individual surface by averaging the values of voxels lying between the gray and white matter surfaces for each surface vertex. For the cerebellum, we computed the non-linear morph into a cerebellar-only version of the Symmetric MNI152NLin2009aSym template (http://nist.mni.mcgill.ca/?p=904). The morph was computed using the SUIT toolbox [107]. The functional data were resampled to a group space of 5446 cerebellar gray-matter voxels with an isotropic resolution of 3mm. During this step, we only considered voxels within the individual cerebellar mask, taking care to exclude any signals from the directly abutting neocortical regions.

### Data types

Starting from the minimally preprocessed time series data, we then compared different ways of filtering, processing, or summarizing these data.

#### Raw Time series

After minimal pre-processing, we calculated covariance matrices based on the resultant functional time series data from either resting-state or the multi-task paradigms.

#### Cleaned time series

We used a single-subject independent component analysis (ICA), implemented in MELODIC (Multivariate Exploratory Linear Optimized Decomposition into Independent Components) and the automated classifier FIX [67–69] to remove structured noise from functional data of the resting-state or task data. Because existing labeled training datasets differed substantially in their study population and acquisition parameters from our data, we created our own training set by manually classifying a subset of runs [67–69]. We selected 17 of the 34 rest runs and 50 of the 272 task runs in the MDTB dataset for hand classification. The cleaned time series were then used as input data to calculate the covariance matrix.

#### Connectivity fingerprint

Starting from the cleaned time series, we conducted an additional processing step that has been shown to be potentially beneficial for subsequent network analysis [70, 71]. To obtain functional connectivity fingerprints from the time series data, we used previously estimated connectivity weight maps [64], consisting of 32 neocortical networks. We then regressed the 32 group network spatial maps into each subject’s data, resulting in 32 subject-specific network time courses. The connectivity fingerprints were calculated as Pearson’s correlations of each voxel time series with each cortical network time course. Covariance matrices (and subsequent parcellation and connectivity models) were then calculated using these 32-dimensional connectivity vectors as input data.

#### Task activation estimates

To estimate task-evoked activation, we used a general linear model (GLM) implemented in SPM [108]. The GLM was fitted to the time series of each voxel separately for each imaging run. We omitted the standard high-pass filtering and instead relied on SPM’s high-dimensional autocorrelation model (FAST) during GLM estimation for temporal filtering, a step that we found to result in more reliable activation estimates across runs [47]. The 5-s instruction phase was modeled with a single regressor but excluded from subsequent analyses. Each task block was modeled with a 30-s boxcar regressor, or with multiple regressors when task blocks contained sub-conditions. Rest was modelled explicitly like all the other tasks. Finally, we subtracted the mean across all task conditions from each of the activation maps, such that all activation estimates were relative to the mean across all tasks. The resultant beta coefficients were univariately pre-whitened by dividing them by the square root of the residual mean-square image, yielding normalized activation estimates for each voxel.

For the HCP task data, a GLM was fitted to the time series of each voxel for each imaging run and session using the FEAT design files provided with the HCP 1200 subjects data 2017 release. As for the other data sets, no spatial smoothing was applied.

#### Task residuals

We derived the residual time series from the task-based data by multiplying the task contrast images with the temporally pre-whitened design matrix (using SPMs high-dimensional autocorrelation model) and subtracting it from the temporally pre-whitened time series.

### Task Battery sampling

To simulate the effect of different task battery sizes on covariance matrix estimation, we created task batteries consisting of 2, 3, 6, 10, 13, or 17 tasks by randomly sampling from the available 17 tasks. To keep the amount of scan time used for each battery the same, we targeted 20 minutes of data for calculating each covariance matrix. Since each task block lasted 30 seconds (0.5 min), we calculated the number of runs needed as the floor of 20*min/*(*n*_tasks_ × 0.5*min*) . This resulted in using 2 runs for 17-task batteries (17 min data), 3 runs for 13-task batteries (19.5 min), 4 runs for 10-task batteries (20 min), 6 runs for 6-task batteries (18 min), 13 runs for 3-task batteries (19.5 min), and 16 runs for 2-task batteries (16 min). The amount of data used for each covariance matrix type was therefore matched as closely as possible given the available data and the constraint that each task block was 30 seconds. For each combination of *n* tasks, we used 30 random task combinations for each subject, resulting in 30 2-task combinations, 30 3-task combinations, and so on.

### Covariance matrices

The spatial (voxel-by-voxel or parcel-by-parcel) covariance matrix is a critical intermediate quantity in many imaging analyses. Most subsequent parcellation [10, 11, 50, 51], gradient [52, 53], and connectivity analyses [39,57 –61] are (implicitly or explicitly) based on this quantity. When it is calculated the original underlying timeseries or activation patterns are no longer required. Thus, the covariance matrix contains all the critical information for these subsequent analyses (i.e. it is a sufficient statistic [63]), and therefore provides an ideal point at which different pre-processing steps can be compared.

For the neocortex, we used a parcellation of each hemisphere into 1442 regular icosahedron parcels. Excluding all parcels that were located within the medial wall, this resulted in 2686 parcels for the entire cortical surface. We then calculated a parcel-by-parcel covariance matrix. For the cerebellum, we calculated a voxel-by-voxel covariance matrix in MNISymC3 space with 5546 voxels. We also calculated a cortico-cerebellar covariance matrix (5546 × 2686, representing the covariance between neocortical parcels and cerebellar voxels.

The covariance matrices were computed from time series data (including residual time series), connectivity fingerprints calculated on that time series data, or from task activation estimates (see above). The covariance matrices were calculated as 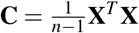, where **X** is the mean-centered data matrix of shape (*n*× *p*), with *n* being the number of observations (conditions or time points) and *p* being the number of parcels or voxels. Mean-centering was performed separately for each run to remove run-specific baseline effects.

### Split-half reliability

We quantified the reliability of covariance matrices based on different data types within subjects by splitting the data into two halves (10 minutes of scan time, a single run), and calculating covariance matrices separately on each half. As a measure of reliability, we then computed the Pearson correlation between the vectorized covariance matrices. We calculated the split-half reliability for each subject and brain structure separately, averaging across voxels to obtain a single reliability measure per subject and data type.

### Cross-tentorial correlation ratio

We quantified the correlations between the most superior cerebellar voxels with abutting neocortical voxels and non-abutting voxels in MNI space to obtain a measure of noise influence on the data. For this, we obtained a mask of the neocortical voxels abutting the cerebellum by dilating the cerebellar mask from the SUIT pipeline [109] by 10 mm. We then projected the voxels that intersected with neocortical grey matter (see red voxels in Figure 4a) onto the fs32k neocortical surface via each subject’s reconstructed pial and white matter surface. We obtained a mask of the superior cerebellar voxels abutting the neocortex by dilating a cerebral mask of the MNI152NLin6Asym wholebrain template by 10 mm and resampling the dilated voxels that intersected with cerebellar grey matter to MNISymC3 space (see blue voxels in Figure 4a).

We then calculated the normalized ratio between superior cerebellar and adjacent neocortical voxel correlations and superior cerebellar voxels and the rest of the neocortex as

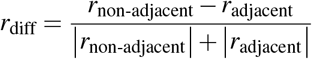

Given the extensive cortico-cerebellar connectivity, especially with more dorsal cortical regions [65, 74], non-abutting voxels are expected to exhibit stronger functional connectivity than abutting voxels. Therefore, we would expect on average positive values of *r*_diff_; smaller or more negative values would indicate an influence of measurement noise.

### Dependence on task set

To examine the dependence of covariance matrices on the specific choice of tasks, we calculated the similarity between different randomly sampled task batteries of the same size. For each task battery size (2, 3, 6, 10, 13, or 17 tasks), we computed pairwise correlations between all covariance matrices derived from the 30 different task combinations. For the 17 task battery the tasks were always the same, but the estimates relied on estimates from separate runs. For each subject and task battery size, we averaged across all pairwise comparisons to obtain a single measure, which quantifies how similar different random task batteries of the same size are to each other.

#### Generalization to novel task set

To evaluate how well covariance matrices derived from different data types generalize to novel task states, we calculated the similarity between each data type’s covariance matrix and a benchmark covariance matrix from the same individual. For the MDTB dataset, the benchmark was a covariance matrix derived from 18 unique task conditions acquired in a separate session that were not included in the training data set (see Supplemental table 1,2). The estimate for the unique task conditions was based on data from all 16 runs, effectively using 144 minutes of functional imaging data).

To approximate how similar the covariance matrices derived from the training set and benchmark data should be if they were truly identical, we correlated the covariance matrices estimated on the activation estimates from 14 runs of the test set with a covariance matrix estimated on the activation estimates from 2 (non-overlapping) runs of the test set. This provides a noise ceiling for interpreting the similarity values, as the similarity is limited by the measurement reliability.

For the HCP dataset, we used a leave-one-task-out approach where the benchmark was a covariance matrix derived from two runs of the task dataset (working memory, gambling, motor, language, social cognition, relational processing, or emotion processing). We estimated the task-activations for each task condition in these runs including the resting baseline. The estimates were then mean-subtracted across conditions in each run and concatenated before spatial covariance estimation.

### Similarity space of covariance matrices

To understand the overall structure of similarity and differences of estimated covariance matrices, we calculated Pearson correlation of covariance matrices, using the different task combinations (see above), the different data types (task and rest time series), as well as the independent task session covariance matrix. The correlation matrix of covariance matrices was then averaged across individuals. To visualize the similarity space, we performed classical multidimensional scaling (MDS) on the mean correlation matrix. Each task battery combination and task type was represented as a point in the resulting low-dimensional space, allowing us to visualize how different sized task batteries cluster based on their similarity, and how they systematically differ from those based on time series data.

### Correlation matrices

While many models rely on the covariance matrix as a sufficient statistic, an alternative choice would have been to look at the spatial correlation matrices instead. Correlation matrices can be derived from the covariance matrices by dividing each row and column by the standard deviation for that voxel or parcel. This normalization removes the influence of variance differences between parcels or voxels, focusing solely on the pattern of co-variation.

We calculated neocortical, cerebellar, and cortico-cerebellar correlation matrices for all data types and repeated all analyses—including split-half reliability, cross-tentorial correlation ratios, task battery similarity, generalization to novel task sets, parcellation, and connectivity modeling—using correlation matrices instead of covariance matrices.

### Parcellations

We derived individual functional parcellations for the multi-task and resting-state data using a two-step approach. Starting from the group atlas for the cerebellum [64] or the neocortex [11], we estimated the individual average time series, connectivity fingerprint, or task activation profile (depending on data type) for each of the parcels, resulting in the average profile **v**_*k*_ for each of the *k* parcels. This was done by fitting a von-Mises-Fisher mixture model to the data of each subject separately using the group atlas as a prior, as described in the hierarchical Bayesian parcellation framework [77]. In the second step, we then correlated the data from each voxel or vertex with each of the averaged profiles from each parcel and assigned the brain location to the one with the highest correlation.

While the hierarchical Bayesian parcellation framework [77] can leverage the advantage of task-activation data by estimating the regional response profiles for an entire group of subjects, we performed the parcellation here individually to ensure a fair comparison with the resting-state data, where the time series for each region will differ across individuals. We also did not apply the spatial group prior in the second step, but assigned the parcels based in the individual data only (given the **v**_*k*_). While the Bayesian integration with a group map has been shown to improve individual parcellations [64, 77], we wanted to evaluate the quality of the individual data types without the biasing influence of a group prior.

#### Evaluation

To assess how well a given parcellation can predict functional boundaries in the cerebellum, we utilized the Distance-Controlled Boundary Coefficient (DCBC) [80]. This metric compares correlations for within-parcel voxel pairs to between-parcel voxel pairs, while accounting for their spatial distance. A higher DCBC value indicates better performance of the parcellation. In contrast to other evaluation metrics (homogeneity or silhouette coefficients), random, but spatially smooth, parcellations have an average DCBC value of zero. The DCBC was calculated using the hard (winner-take-all) individual parcellation to predict the functional boundaries in held-out data of the same individual.

### Connectivity modeling

To examine the connectivity from neocortex to cerebellum we used an L2-regularized regression (Ridge regression) model [65]. Each model was trained and evaluated on individual subjects separately.

The neocortical data were averaged with regular Icosahedron parcels (see covariance matrix), resulting in a *N* × 2686 matrix **X**_*i*_ (neocortical activation of subject *i*). For the cerebellum we used all voxels in our template resulting in the *N* × 5446 matrix **Y**_*i*_ (cerebellar activation of subject *i*). Using ridge regression, we then estimated the matrix of connectivity weights:

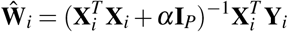

As can be seen from this equation, the connectivity model is based on the sufficient statistics of the neocortical 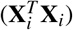 and cortical-cerebellar 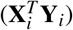 covariance matrices. We used a wide range of regularization coefficients log(*α*)= *{*2,…, 14*}* for each datatype.

#### Evaluation

All models were evaluated on the task activation estimates for the unique tasks from Task set B of the MDTB dataset. For each model, the regularization term that led to best performance across participants was chosen. Using the connectivity weights from the training set (**Ŵ** ) and the neocortical data from test set (**X**_test_), the cerebellar data from the test set (**Y**_test_) were predicted using **Ŷ** _test_ = **X**_test_**Ŵ** . The correlation of **Y**_test_ and **Ŷ** _test_ was reported as the evaluation metric.

To prevent the model evaluation being biased by spatially structured measurement noise, the activation estimates of the test data were estimated on each half (8 runs) of the data separately. The cerebellar data from the second half was then predicted using the cortical data from the first half, and vice versa. Reported evaluation results are averaged across the two halves.

### Statistical analysis

Means are reported ± the standard error of the mean across individuals. Unless otherwise noted, we report paired t-tests, treating individuals as the random effect. We always report exact uncorrected p-values in the text, but note if these do not pass a Bonferroni-corrected threshold, adjusted for the number of tested data types, if appropriate.

## 1 DATA AVAILABILITY

The raw task and rest fMRI data of the MDTB has been deposited in the openneuro database under accession code ds002105 [doi:10.18112/openneuro.ds002105.v1.1.0]. For the HCP dataset, raw and preprocessed data is available at https://www.humanconnectome.org/study/hcp-young-adult/data-releases. The replication dataset has not yet been openly released.

## 2 CODE AVAILABILITY

The code for generating the results and figures in this paper is publicly available as the GitHub repository https://github.com/carobellum/RestMultitask.git. The code for the hierarchical Bayesian parcellation framework is available at https://github.com/DiedrichsenLab/HierarchBayesParcel. The organization, file system, and code for managing the diverse set of datasets is available at https://github.com/DiedrichsenLab/Functional_Fusion. The code for connectivity modelling is available at https://github.com/DiedrichsenLab/cortico_cereb_connectivity. The Python toolbox for running multi-domain task batteries, along with an implementation of most of the tasks used in the current batteries is openly available at https://github.com/diedrichsenlab/MultiTaskBattery.git.

## ACKNOWLEDGMENTS

This work was supported by a Discovery Grant from the Natural Sciences and Engineering Research Council of Canada (NSERC, RGPIN-2016-04890), and a project grant from the Canadian Institutes of Health Research (CIHR, PJT-191815), both to J.D. Additional funding came from the Canada First Research Excellence Fund (BrainsCAN) to Western University. C.R.N. was supported by a Wellcome Trust Early Career Award (306553/Z/23/Z) and a Junior Research Fellowship Grant from Linacre College, University of Oxford.

## AUTHOR CONTRIBUTIONS

C.N. and J.D. conceived the study, J.D. guided the study design, C.N., B.A. and M.S. collected the data, J.D. guided the analysis, J.X. and A.L.P. provided data analysis tools, C.N. and A.S. performed the analysis, C.N. and J.D. wrote the manuscript, all authors reviewed and approved the manuscript.

