## Supplementary Materials for "Multi-task fMRI outperforms resting-state fMRI for revealing task-invariant organization of the human brain"

#### S1. Reliability of cerebellar and cortico-cerebellar covariance matrices

Cerebellar and cortico-cerebellar covariance matrices estimated on the task-evoked activation showed substantially lower split-half reliability than all other data types with the exception of raw task and rest time series and, in the case of cerebellar covariance, residual time series (all $t_{16} < -3.14, p < 6.64 \times 10^{-3}$; significant at Bonferroni-adjusted threshold of $p < 0.007$).

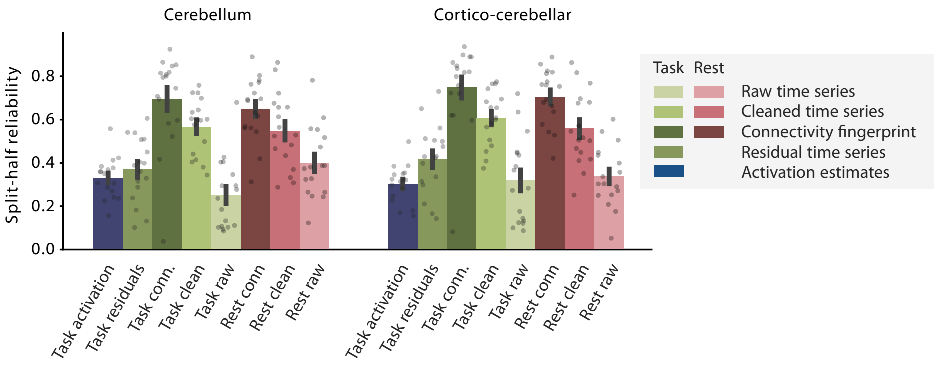


**Figure S1. Split-half reliability for cerebellum and cortico-cerebellar covariance matrices.** Split-half reliability was calculated as the correlation between the covariance matrices derived from two halves of the cerebellar or cortico-cerebellar data. *Raw time series*: raw fMRI time series without further processing. *Cleaned time series*: fMRI time series after ICA-based denoising. *Conn. fingerprint*: Connectivity fingerprint calculated on cleaned time series data. *Task activation estimates*: GLM-modelled task data of 17 tasks. *Residual time series*: Task time series where the task-related activation has been regressed out. Error bars represent standard error of the mean across participants. Each dot is one participant.

#### S2. Dependence on exact choice of task battery for cerebellar and cortico-cerebellar covariance

Inter-battery similarity increased significantly from 3-task to 6-task batteries for both cerebellum and cortico-cerebellar covariance (all $t_{16} = -2.73$, $p < 1.50 \times 10^{-2}$). For both cerebellar and cortico-cerebellar covariance, the similarity between different task batteries also reached its plateau at 6 tasks.

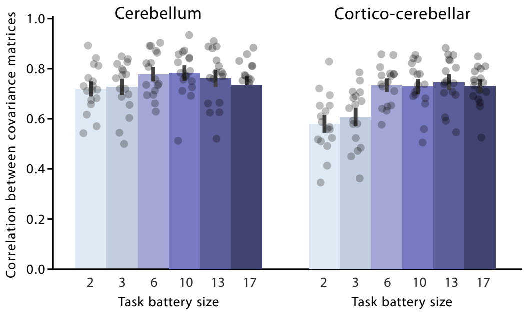


**Figure S2. Correlation of cerebellar and cortico-cerebellar covariances matrices estimated on different batteries**. Pairwise correlations between the covariance matrices from cerebellar or cortico-cerebellar data of all task combinations. Error bars represent standard error of the mean across participants. Each dot is one participant.

### S3. Generalization to novel task set for cerebellar and cortico-cerebellar covariance matrices

As for the neocortex, task activation estimates from 17 tasks outperformed all other data types (cerebellum: all $t_{16} > 5.50, p = 3.37 \times 10^{-4}$ significant at Bonferroni-adjusted threshold of $p < 0.007$, cortico-cerebellar: all $t_{16} > 2.33, p < 3.33 \times 10^{-2}$) apart from the task connectivity fingerprint (cortico-cerebellar: $t_{16} = -1.23, p = 0.23$, cerebellar: $t_{16} = 0.43, p = 0.67$) and, for cerebellar covariance, the cleaned task time series (task activation vs. cleaned task time series: $t_{16} = -0.06, p = 0.95$). Task activation estimates from 17 tasks also outperformed task batteries 2 tasks (cerebellum: $t_{16} = 4.14, p = 7.73 \times 10^{-4}$, cortico-cerebellar: $t_{16} = 9.84, p = 3.43 \times 10^{-8}$) and, in the case of cortico-cerebellar activation, 3 tasks ($t_{16} = 5.48, p = 5.06 \times 10^{-5}$). We did not observe a significant difference to batteries with 6 tasks ($t_{16} = 2.01, p = 6.14 \times 10^{-2}$) or 10 tasks for cortico-cerebellar covariance ($t_{16} = -1.47, p = 1.61 \times 10^{-1}$), and 13 task batteries showed significantly higher similarity than 17 tasks ($t_{16} = -4.16, p = 7.37 \times 10^{-4}$). Further, for cerebellar covariance, the task activation estimates of 6, 10 and 13 tasks showed higher similarity with the new tasks than 17 tasks (6 tasks: $t_{16} = -2.36, p = 3.13 \times 10^{-2}$, 10 tasks: $t_{16} = -4.21, p = 6.65 \times 10^{-4}$, 13 tasks: $t_{16} = -2.82, p = 1.22 \times 10^{-2}$). This pattern may reflect a trade-off between task diversity and per-task data quantity: increasing the number of tasks reduces the amount of data available per task, which may disproportionately affect estimates that involve the cerebellum given its lower signal-to-noise ratio.

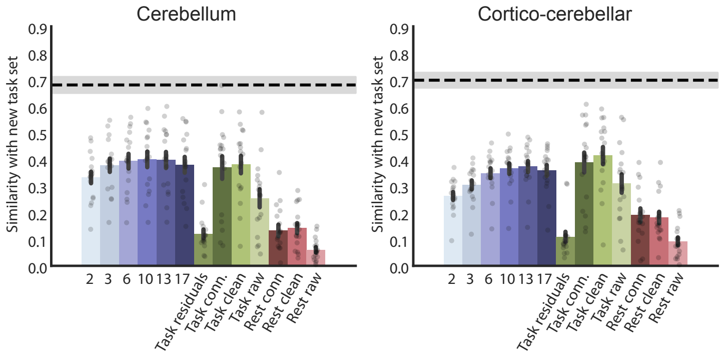


**Figure S3. Similarity for cerebellum and cortico-cerebellar covariance matrices to independent task set**. Similarity is the correlation of covariance matrices with the covariance matrix calculated from the same individual, but using 18 tasks novel tasks, acquired on a separate imaging session. Noise ceiling was calculated as split-half reliability of the independent task session on which all similarities were evaluated. Error bars represent standard error of the mean across participants. Each dot is one participant.

### S4. Generalization to novel task set for correlation matrices

We repeated the results shown in Figure 6 using correlation, rather than covariance matrices. Again, task time series outperformed rest time series for matched preprocessing steps, including the raw time series (cortex: $t_{16} = 4.16, p = 7.42 \times 10^{-4}$; cerebellum: $t_{16} = 7.28, p = 1.85 \times 10^{-6}$; cortico-cerebellar: $t_{16} = 8.37, p = 3.07 \times 10^{-7}$), the cleaned time series (cortex: $t_{16} = 2.44, p = 2.68 \times 10^{-2}$; cerebellum: $t_{16} = 7.06, p = 2.69 \times 10^{-6}$ ; cortico-cerebellar: $t_{16} = 7.58, p = 1.11 \times 10^{-6}$) and as connectivity fingerprints (cortex: $t_{16} = 2.15, p = 4.75 \times 10^{-2}$; cerebellum: $t_{16} = 9.06, p = 1.06 \times 10^{-7}$; cortico-cerebellar: $t_{16} = 8.37, p = 3.05 \times 10^{-7}$).

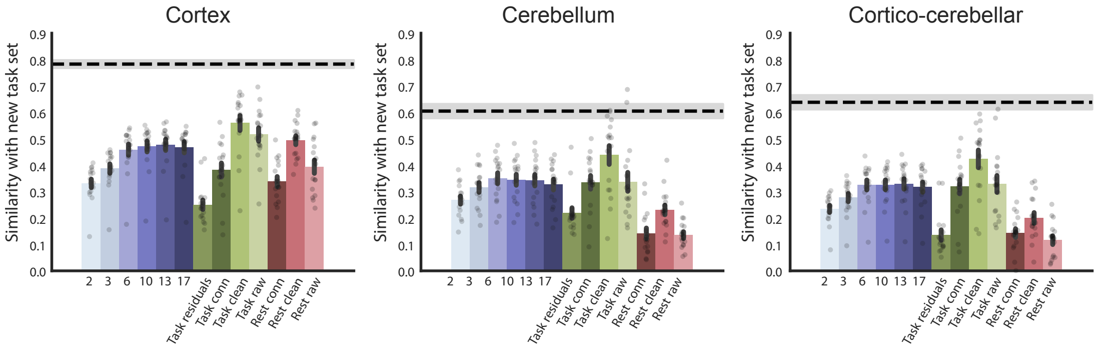


**Figure S4. Similarity for cortical, cerebellar and cortico-cerebellar correlation matrices**. Similarity is calculated as correlation of correlation matrices with the same individual's new task session. Noise ceiling was calculated as split-half reliability of the independent task session on which all similarities were evaluated. Error bars represent standard error of the mean across participants. Each dot is one participant.

### S5. Replication in the Human Connectome Project (HCP)

We replicated our results shown in Figure 6 on a commonly used dataset in the field, the task and rest data from the Human Connectome Project (HCP). We tested how well task data and rest data predicted the spatial covariance matrix based on the two runs of a single task (working memory, gambling, motor, language, social cognition, relational processing or emotion processing). For task data, we used task activation estimates, based on 14 task-based runs, each time leaving out the task runs that were used for evaluation. With each task run lasting between 2.3 to 5 minutes, each spatial covariance estimate was therefore based on 38.8 - 47.6 minutes. For rest data, we used four imaging runs of the cleaned time series, each run lasting 15 minutes. Therefore the spatial covariance matrix for the rest data was based on 60 minutes of imaging data. We evaluated these estimates by correlating them with a spatial covariance matrix based on the task activation estimates of the 2 left-out runs (see Supplementary Figure S5). The results were averaged across cross-validation folds to arrive at one value per participant.

The similarity to the unrelated tasks was significantly higher for the task data than for the rest data for cortical ($t(49) = 6.13, p = 1.50 \times 10^{-7}$), cerebellar ($t(49) = 2.76, p = 8.03 \times 10^{-3}$), and cortico-cerebellar ($t(49) = 4.82, p = 1.43 \times 10^{-5}$) covariance matrices.

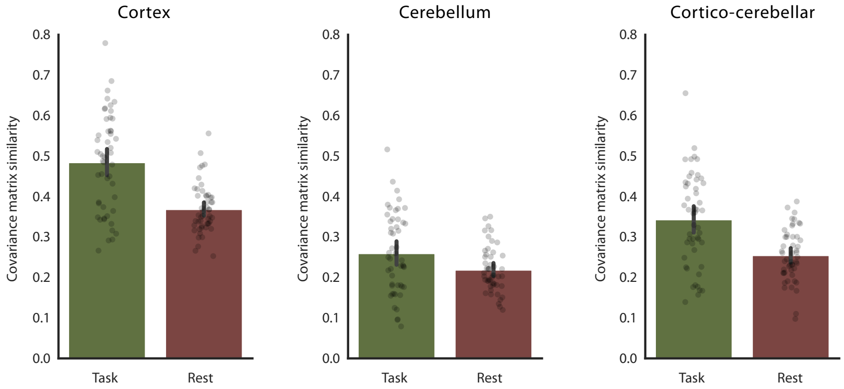


**Figure S5. Similarity for cortical, cerebellar and cortico-cerebellar covariance matrices in the Human Connectome Project (HCP).** Similarity was calculated as the Pearson correlation between covariance matrices derived from all tasks except one held-out task and the covariance matrix of the held-out task (leave-one-task-out cross-validation). For task data, covariance matrices were computed from task activation estimates (two runs per task; 38.8 - 47.6 min total). For resting-state data, covariance matrices were computed from cleaned time series (four runs; 60 min total). Results were averaged across cross-validation folds. Error bars represent the standard error of the mean across participants (N = 50). Each dot represents one participant.

### S7. Parcellation results for prefrontal cortex

In the prefrontal cortex, cleaning and connectivity fingerprinting of the time series data increased performance of parcellations compared to the raw signal (cleaning: $t_{16} = 3.55, p = 5.28 \times 10^{-3}$; fingerprinting: $t_{16} = 7.15, p = 4.61 \times 10^{-6}$) and cleaning, but not fingerprinting improved performance for the rest data (cleaning: $t_{16} = 13.70, p = 5.92 \times 10^{-10}$; fingerprinting: $t_{16} = 2.29, p = 0.072$).

### S8. Parcellation results for cerebellum

In the cerebellum, task data outperformed rest data for matched preprocessing (Raw time series: $t_{16} = 7.13, p = 7.22 \times 10^{-6}$; cleaned time series: $t_{16} = 4.77, p = 6.23 \times 10^{-4}$; connectivity fingerprint: $t_{16} = 8.49, p = 7.61 \times 10^{-7}$). Cleaning increased performance of task data ($t_{16} = 4.23, p = 1.27 \times 10^{-3}$), but fingerprinting did not ($t_{16} = -0.06, p = 1$). Meanwhile, cleaning improved the performance of rest data ($t_{16} = 6.04, p = 3.44 \times 10^{-5}$) while fingerprinting decreased it in the cerebellum ($t_{16} = -5.29, p = 1.47 \times 10^{-4}$).

### S9. Parcellation results for replication dataset

We used a separate Multi-task dataset from our laboratory as a replication dataset (see methods). All models were trained on 40 minutes of imaging data (runs 1-4 of the localizer session for task-based models, or 40 minutes of rest for resting-state models). The evaluation was performed on 40 minutes of held-out multi-task fMRI (runs 5-8 of the same localizer session) using the Distance-Controlled Boundary Coefficient (DCBC) to measure how well each parcellation predicted functional boundaries in these novel task data.

As expected, cleaning resulted in improved parcellations for rest ($t_7 = 3.09, p = 3.49 \times 10^{-2}$) but not task data ($t_7 = 1.73, p = 0.253$), whereas fingerprinting did not improve the resulting parcellations in these eight subjects (all $t_7 < -2.30, p > 0.11$). Parcellations derived from task activation estimates showed highest performance and significantly outperformed task connectivity fingerprint ($t_7 = 3.56$, $p = 9.20 \times 10^{-3}$), cleaned task time series ($t_7 = 5.56$, $p = 8.53 \times 10^{-4}$), raw task time series ($t_7 = 7.04$, $p = 2.05 \times 10^{-4}$), rest connectivity fingerprint ($t_7 = 8.19$, $p = 7.84 \times 10^{-5}$) and the cleaned and raw rest time series (cleaned: $t_7 = 6.55$, $p = 3.19 \times 10^{-4}$, raw: $t_7 = 10.08, p = 2.03 \times 10^{-5}$). For matched preprocessing, task data significantly outperformed rest data (raw time series: $t_7 = 3.75$, $p = 7.16 \times 10^{-3}$; cleaned time series: $t_7 = 3.31$, $p = 1.30 \times 10^{-2}$; connectivity fingerprint: $t_7 = 5.35$, $p = 1.06 \times 10^{-3}$).

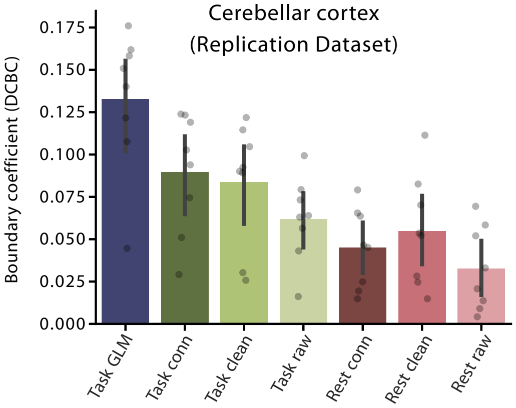


**Figure S6. Parcellation Replication**. Performance of individual parcellation maps in predicting functional boundaries in the cerebellar cortex for the same subjects in the replication dataset. The Distance controlled boundary coefficient (DCBC) indicates the difference of the within-parcel vs. between-parcel correlation, controlled for the spatial distance of two brain locations. For random, but spatially contiguous parcellations, the expected DCBC is zero.

### Supplementary Tables

**Table 1.** **MDTB dataset task set A.**

| Task | Description | Conditions |
|---|---|---|
| Object Viewing | Passive viewing of pictures of objects and a checkerboard pattern | – |
| Motor Imagery | Imagine playing a game of tennis | – |
| Stroop | 3AFC color-naming task with congruent and incongruent color–word mappings | Congruent; Incongruent |
| Verbal Working Memory | 2AFC letter task indicating whether the current stimulus matches the one presented two items earlier | 0-back; 2-back |
| Interval Timing | 2AFC indicating whether a tone is short (100 ms) or long (175 ms) | – |
| Arithmetic | 2AFC judging correctness of simple multiplication equations; control digit judgment task | Math; Digit judgment |
| IAPS Affective | 2AFC indicating whether images are pleasant or unpleasant | Pleasant scenes; Unpleasant scenes |
| IAPS Emotion | 2AFC indicating whether a face depicts happiness or sadness | Happy faces; Sad faces |
| Go/No-Go | Word-based Go/No-Go task with positive (Go) and negative (No-Go) stimuli | Go; No-Go |
| Theory of Mind | 2AFC indicating whether short stories contain true or false beliefs | – |
| Rest | Passive viewing of a fixation cross | – |
| Object N-back | Object-based variant of the letter n-back task | 0-back; 2-back |
| Verb Generation | Covert verb generation or word repetition in response to visually presented nouns | Generate; Read |
| Spatial Imagery | Imagine navigating between rooms in one's childhood home | – |
| Motor Sequence | Execution of a six-element finger sequence or repeated single-finger presses | Finger sequence; Finger simple |
| Action Observation | Passive viewing of videos of knots being tied, with later recall | Video actions; Video knots |
| Visual Search | 2AFC identifying presence of a target among distractors with varying set sizes | Small (4); Medium (8); Large (12) |

**Table 2.** **MDTB dataset task Set B.**

| Task | Description | Conditions |
|---|---|---|
| Spatial Map | Memorize spatial mappings of numbers for later recall | Easy (1); Medium (4); Hard (7) |
| Mental Rotation | Mental rotation task with difficulty defined by angular disparity between target and baseline objects | Easy (0); Medium (50); Hard (150) |
| Biological Motion | 2AFC identifying intact or scrambled point-light walkers | Biological motion; Scrambled motion |
| Concrete Permuted Rules (CPRO) | Apply logic, sensory, and motor rules to sequentially presented stimuli | – |
| Word Prediction | 2AFC judging semantic coherence of word sequences | Prediction; Prediction violated; Prediction scrambled |
| Response Alternatives | Rapid motor response to an imperative signal appearing at varying locations | Easy (1); Medium (2); Hard (4) |
| Nature Movie | Passive viewing of a nature clip from *Planet Earth II: Islands* | – |
| Animated Movie | Passive viewing of an emotional clip from the Pixar movie *Up* | – |
| Landscape Movie | Passive viewing of an aesthetically pleasing landscape clip | – |

**Table 3.** **Replication dataset tasks.**

| Task | Description | Conditions |
|---|---|---|
| Rest | Passive viewing of a fixation cross | – |
| Action Observation | Passive viewing of videos of knots being tied, with later recall | Video actions; Video knots |
| Verb Generation | Covert verb generation or word repetition in response to visually presented nouns | Generate; Read |
| Intact Passage | Listen to clearly spoken sentences [110] | – |
| Degraded Passage | Listen to acoustically degraded sentences [110] | – |
| Sentence Reading | Read sentences presented word by word [111] | – |
| Nonword Reading | Read pronounceable nonwords [111] | – |
| Auditory Narrative | Listen to an extended spoken narrative (Moth Radio Hour) | – |
| Theory of Mind | 2AFC indicating whether short stories contain true or false beliefs | – |
| Tongue Movement | Alternate tongue movements from right to left at 1 Hz | – |
| Motor Sequence | Execution of a six-element finger sequence | – |
| Animated Movie | Passive viewing of an emotional clip from the Pixar movie *Up* | – |
| Object N-back | Object-based variant of the n-back task, indicating whether the current stimulus matches the one presented two items earlier | – |
| Oddball | Detect a target stimulus among distractors | – |
| Demand Grid | Solve complex spatial reasoning problems [112] | – |
| Spatial Imagery | Imagine navigating between rooms in one's childhood home | – |
